# Epileptiform Discharges Drive Unique High-frequency Oscillations Within the Retrosplenial Cortex of Mice with Third Trimester Alcohol Exposure

**DOI:** 10.64898/2026.06.04.729968

**Authors:** Abbey Myrick, Sam McKenzie, Fernando Valenzuela, David Linsenbardt

## Abstract

Fetal Alcohol Spectrum Disorders (FASDs) are associated with alterations in learning and memory that persist throughout the lifespan. Thus, determining the neural mechanisms driving these alterations has the potential to identify novel therapeutic targets for improving memory in those with FASD. Given the newly realized role of the Retrosplenial cortex (RSC) for learning and memory, as well as the profound neural apoptosis that exposure to alcohol during development causes to this brain region, we recorded electrophysiological activity from mice exposed to alcohol during the third trimester-equivalent developmental time period. We observed a large number of Epileptiform Discharges (EDs) in alcohol-exposed subjects compared to controls, which were found to drive with High-frequency Oscillations (HFOs). Furthermore, many features of HFOs (amplitude/duration/etc.) were found to be directly proportional to the temporal distance from ED onset. These findings identify EDs for the first time as a critical feature in a preclinical model of FASD, and suggest their relationship to RSC HFOs may be a key mechanism driving memory alterations.

## Introduction

Learning and memory differences among individuals living with Fetal Alcohol Spectrum Disorder (FASD) are well established and represent significant hurdles to daily life. Among the many brain regions regulating these learning and memory alterations, the retrosplenial cortex (RSC) has emerged as a both critical for memory function and profoundly impacted by developmental alcohol exposure^1–8^. Specifically, the RSC receives hippocampal Sharp-Wave Ripples via the Subiculum—high frequency oscillations (HFOs) hypothesized to move new memories out to further cortical regions from their inception in the hippocampus, making the retrosplenial cortex a gateway for memory consolidation^1,2,5,9–11^. Despite this critical role, the impact of developmental alcohol-induced damage to HFOs within the RSC has not been directly investigated.

Epileptiform discharges (EDs) are large amplitude electrical events that have been documented as not only occurring co-morbid with epilepsy, but also without seizures, in Alzheimer’s disease, Parkinson’s disease and following brain injury^12–18^. EDs have been shown to negatively impact memory and are hypothesized to partially underlie memory disruptions in these conditions^13,19,20^. Because of this, EDs are becoming recognized as a symptom of network dysfunction, developing from neuromodulation — such as neurodevelopmental disorders may cause. Importantly, compelling clinical research has shown elevated risk of abnormal electroencephalogram (EEG) readouts, including EDs, and the development of epilepsy among individuals with FASD^17,21–24^. Though statistical power is often an issue with clinical studies of this population, the estimated risk of atypical EEG at 3-21% should motivate researchers to pursue investigating EEG events further.

Preclinical research on potential mechanisms for this elevated risk of atypical EEG are sparse. Here, we characterize EDs in the retrosplenial cortex and investigated their potential role in driving HFOs and thus, memory function. We discovered many more EDs within our alcohol treated subjects, with a subset closely followed by HFOs which were greater in power, frequency, duration and amplitude compared to HFOs falling outside of EDs. Further research on the link between epileptiform activity and FASD is a logical step in teasing apart the mechanisms behind memory differences and the current study provides a preclinical model to do so.

## Methods

### Animals

C57BL/6J mice were acquired from JAX laboratory and bred in house. Third trimester-equivalent alcohol exposure was modeled using two doses of 20% alcohol in saline solution (2.5 g/kg) spaced two hours apart and delivered subcutaneously at postnatal day 7 (n= 5) while saline controls received saline injections (n=5). Following physical maturation at postnatal day ∼60, mice were anesthetized with isoflurane and surgically implanted with Cambridge Neurotech silicone single shank L3 series probes painted with the fluorescent red dye Dil (Sigma, St. Louis, MO, USA) with Intan 64-channel head stages at the stereotaxic coordinates in the retrosplenial cortex (AP = -1.79 to -2.02, ML = 0.52, DV = 3.0). Mice were single housed in 6×10×5 inch amber cages, and provided peanut chips and gel nutrition until fully recovered from surgery. All animals were kept in a reverse 12/12 light/dark cycle with lights going off at 9am and coming on at 9pm. 24-hour recordings were switched over between 1-3pm every day. *Ad lib* food (PicoLab Laboratory Rodent Diet 5LOD) and water (treated by the New Mexico Animal Resource Facility). At the conclusion of the study, animals were euthanized using Ketamine injection until no response to an eye touch was present, then were perfused with Phosphate Buffered Saline (PBS) and paraformaldehyde (PFA). Brains were removed, and drop fixed in 30% sucrose before transferring to PBS and stored in -20 before slicing on a cryostat. Sections were stained for DAPI (4′,6-diamidino-2-phenylindole) and mounted on Superfrost Plus Gold (Fisherbrand) slides with 22 × 50 mm coverslips and Vectashield (Vector Laboratories) mounting media and probe placement was visually verified where possible. All procedures were approved by the University of New Mexico Animal Care and Use Committee and conformed to the Guidelines for the Care and Use of Mammals in Neuroscience and Behavioral Research (National Research Council, 2003).

### Data Acquisition

Probe placement was verified during recording via oscillatory markers of granular layer of the retrosplenial cortex and the presence of hippocampal activity when available. When needed, probe depth was adjusted and tissue allowed to settle for 24-hours before collecting data. All recordings were done in the animal’s home cage using the OpenEphys acquisition system at a 30,000 Hz sampling rate. Commutators (Doric: order code AERJ_24_) allowed for free movement of the animal during all recordings. Data were down-sampled to 1250 Hz and analyzed using custom MATLAB code with Neuroscope2 (CellExplorer) for visualization.

Detection for EDs was run using original MATLAB code. These timestamps were further refined by locating the absolute largest deflection within +/-15 ms of the original timestamp and setting this as time 0. Power and amplitude metrics were computed using the raw down-sampled signal within a +/-30 ms window around the refined timestamp. For ED cluster analysis, waveforms were reduced from 1250D to 2D space using the *run_umap* function followed by clustering using *kmeans* and a K-value was determined by *evalclusters*.

A +/-10 second window around the ED timestamps was bandpass filtered at 100-250 Hz using a zero-phase 2nd-order Butterworth filter. A 10 ms sliding RMS envelope was computed and normalized to z-scores relative to the mean and standard deviation of the 20 second segment. HFOs were defined as continuous epochs exceeding 2 standard deviations above the mean and were retained if they had a duration of 20-150 ms and met a 3-cycle minimum. HFOs within 20 ms of EDs were excluded as noise and ripples within 5 ms of each other were merged. For visualization, waveforms were aligned to their most negative trough.

### Statistics

All analyses were conducted in MATLAB R2024b and statistical significance was defined as p<05. Wilcoxon rank-sum and signed-rank tests were run on count data using the *ranksum* and *signrank* functions, respectively. All other analyses were done with mixed-effects models. Outcome variables were assessed for normality using the Jarque-Bera (*jbtest*) test and skewness. Continuous variables with positive skewness that were bounded at zero were analyzed using *fitglme* with a gamma distribution and identity link (with prior check for confounding zero values) unless otherwise stated. ED minimum amplitude was log transformed using absolute values prior to analysis with a linear mixed-effects model. Animal ID nested inside day improved model fit across analysis and was included as a random intercept in all models. HFO timestamp distribution statistics were run on absolute values with a before vs after condition. The limited number of EDs in the saline group posed a problem in statistically comparing across treatments and should be considered a limitation where it was performed.

## Results

Following surgical implantation in the retrosplenial cortex (Fig.1a) and recovery, two 24-hour home-cage recordings revealed EDs spanning the entire region with a significant treatment difference in number of EDs (U = 140, p = 0.0101; Fig.1b). While we detected these events in saline mice, the number was extremely low as evidenced by the raw traces in Fig. 1c compared to Fig. 1d. A generalized linear mixed-effects model with log link found no difference in mean maximum amplitude (p = 0.38917) between saline (1008.04 uV; Fig. 1c) and TTAE (907.68 uV; Fig. 1d) though between-animal/day variability was high (SD = 0.3273 log-uV). Likewise, a linear mixed-effects model found no difference in average minimum amplitude (p = 0.4204) between saline (−1160.17 uV) and TTAE (−993.55 uV) with even higher between-animal variability (SD = 0.4325 log-uV). To investigate the possibility of neuroinflammatory responses as the driver behind the extreme number of EDs in our TTAE group, we performed a chronic electroencephalogram (EEG) study to test non-invasively for EDs, which we found occurring on the neocortex with similar frequency as in the deeper silicone probe recordings (Fig. 1b; grey triangles). While these data are displayed alongside the rest of the TTAE group, they were not included in the statistical analysis.

**Fig. 1.**
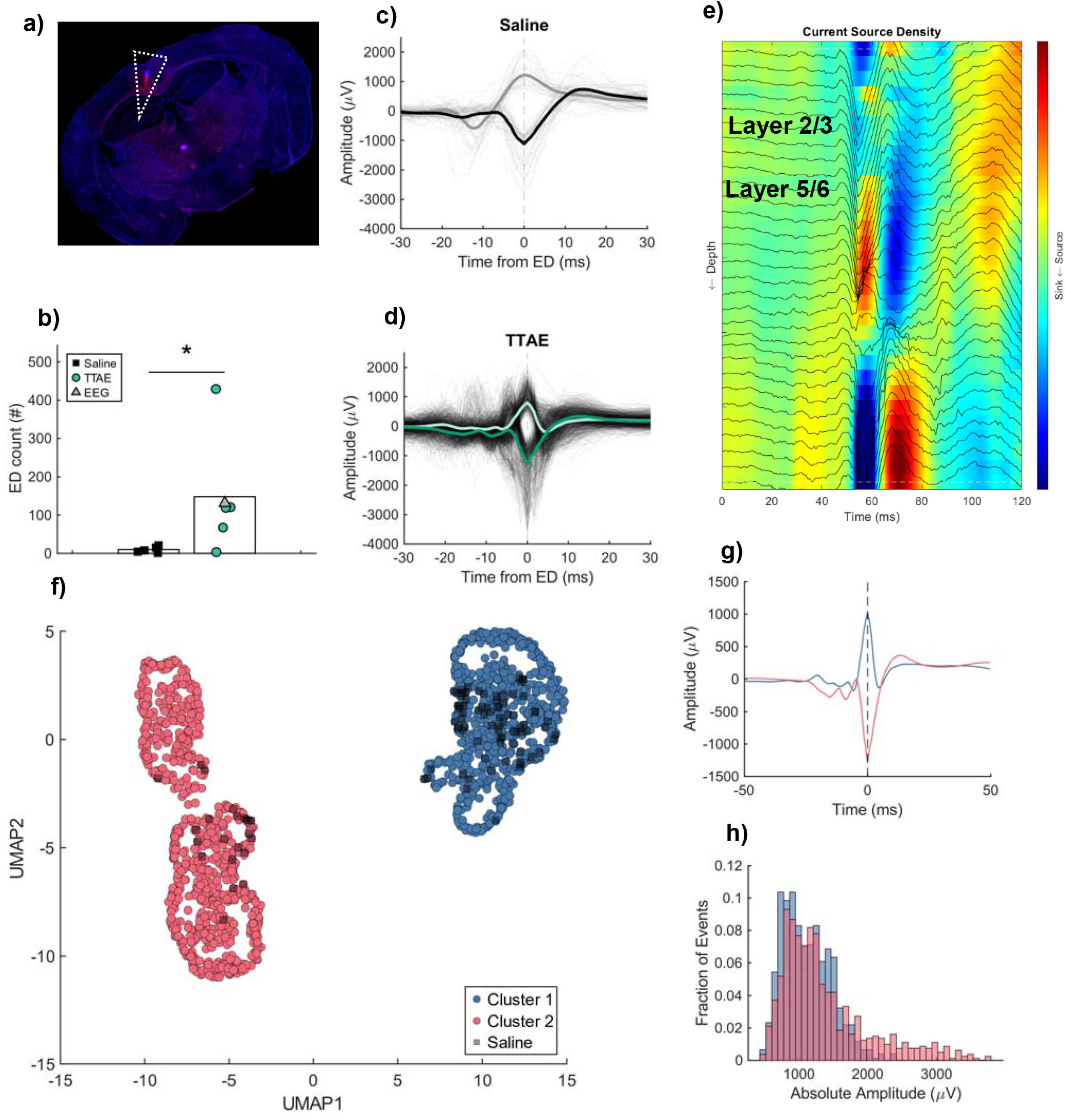
TTAE results in two types of EDs that are mainly characterized by polarity. a) Probe placement spanning the granular layers of the retrosplenial cortex. b) Average number of EDs was significantly different between saline (black square, n = 100) and TTAE (teal circle, n = 1479) (p = 0.012). Grey triangles represent TTAE data collected by EEG (n = 1 mouse). c) Average waveform in the saline group, n = 5 mice; positive = 37, negative = 63, mean. d) Average waveform in the TTAE group, n = 5 mice; positive = 799, negative = 680. e) Example current source density with local field potential plotted on top. f) K-means clustering projected onto reduced feature space. Saline animals are indicated in black (grey?) squares. g) Average waveforms; n = 772 (cluster 1), n = 807 (cluster 2). h) Distribution of ED absolute amplitudes was not significantly different (p = 0.3243).

K-means clustering identified two distinct populations of ED waveforms (Fig. 1f; saline EDs projected in black squares). Cluster 1 and 2 appear to differ predominantly on polarity (Sup.1). Polarity differences are not the result of differences in electrode placement as is apparent in the distribution of both cluster types across each animal (Sup. 2). Though cluster 1 exhibited a larger spread in absolute amplitude, it was not statistically significant (p = 0.3243; 1116.0±352.7 μV, vs 1366.5±650.4 μV, Fig. 1h).

We then investigated effects on HFOs within a 10 second window before and after the EDs. Due to the very low number of events in the saline group, that dataset was not included in the following analysis. In the alcohol exposed group, there was not a significant difference in HFO count pre-vs post-ED (Pre = 1092, post = 1610; U = 114, p = 0.5452) but a significant difference in the *distribution* of timestamps pre-vs post-(β = -1457.2 ms, 95% CI -1741.7, - 1172.6], t(2700) = -10.042; p<0.0001). HFO onset distribution relative to ED onset (Fig. 2a) revealed a large number of HFOs occurring 20-100 ms (mean = 65.7 +/-0.07) after the ED. These ‘Rapid HFOs’ (rHFOs; Fig. 2b, N = 482) exhibited multiple characteristics that were temporally affected by time from the ED. Power decreased with increased time from ED (β = - 244.21 uV, 95% CI [-312.05, -176.37], t(480) = -7.0734, p<0.0001). Time quadratically affected rHFO frequency (β = -6669.46 Hz/_s_^2^, 95% CI [-10768, -2570.5], t(479) = -3.1971, p<0.0015). Duration decreased linearly with time from ED (β = −193.59 ms, 95% CI [−268.92, −118.25], t(479) = -5.0492, p<0.0001). Similarly, maximum amplitude decreased with time from ED (β = - 547.98 uV, 95% CI -710.91, -385.06], t(480) = -6.6088, p<0.0001). ED cluster ID (Fig.1f-g) did not predict the distribution of HFOs and rHFOs within the 20 second window centered on EDs (p = 0.53181), nor did ED cluster ID predict the presence of an rHFO (p = 0.92317).

**Fig. 2.**
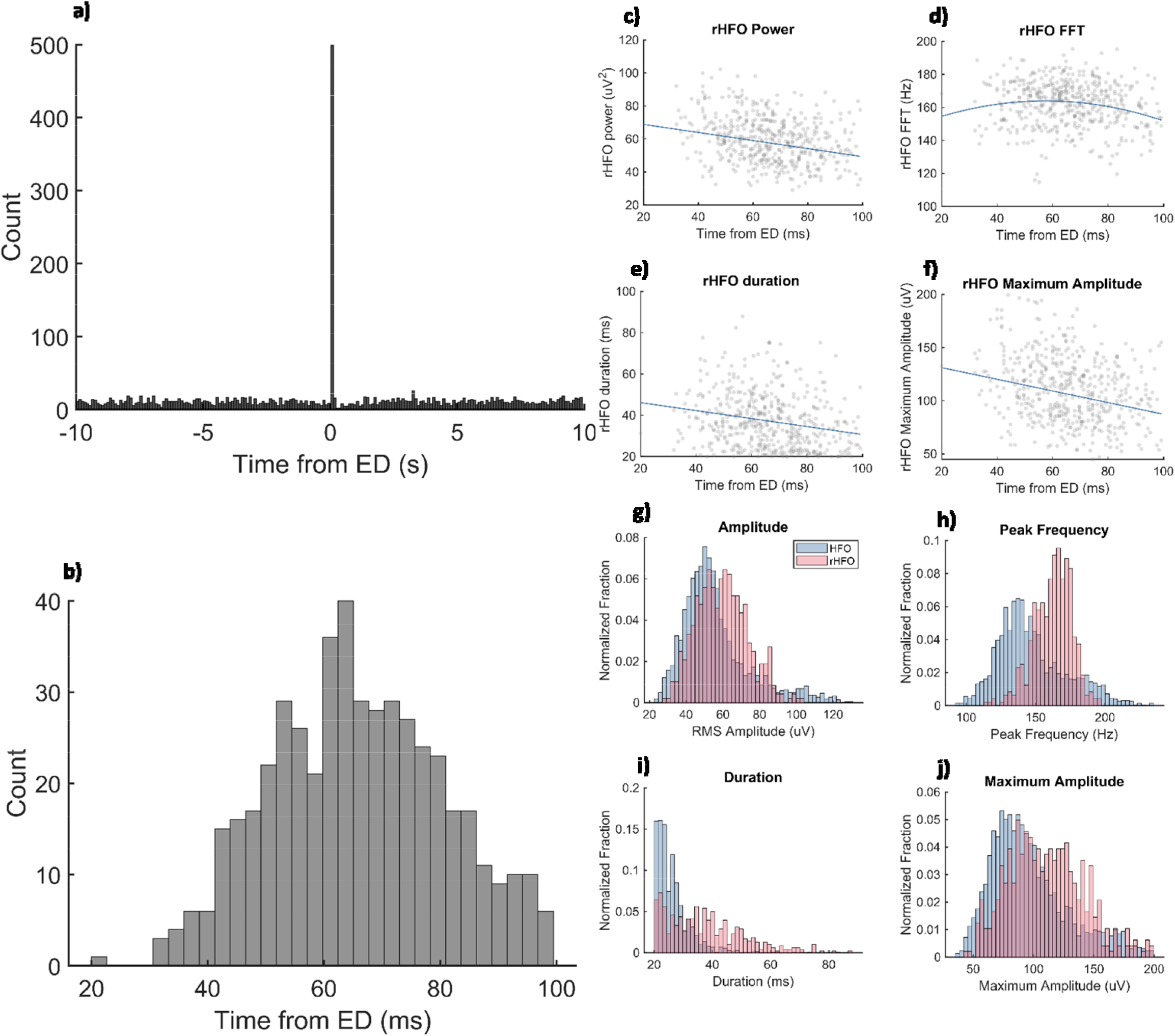
rHFOs in TTAE exhibit altered metrics compared to HFOs. a) HFO onset distribution uncovered a large number of events 20-100 ms after the ED (rHFOs). b) Mean rHFO onset of 65.7 +/-0.07 ms following ED peak absolute amplitude. c) rHFO power decreased with time from ED (p<0.0001). d) rHFO frequency exhibited a quadratic relationship with time from ED (p = 0.0015). e) rHFO duration decreased with time from ED (p<0.0001). f) Maximum amplitude decreased with time from ED (p< 0.0001). g) rHFOs had larger amplitude compared to HFOs (p<0.0001). h) Peak frequency was higher in rHFOs (p<0.0001). i) Duration was longer in rHFOs (p<0.0001). j) Maximum amplitude was higher in rHFOs (p<0.0001).

Previous literature indicates EDs reduce Sharp-Wave Ripple rate in the hippocampus^13,19^ which are known to travel from the hippocampus to the retrosplenial cortex as HFOs via the subiculum, however, not all HFOs in the retrosplenial cortex are sourced from Hippocampal Sharp-Wave Ripples^5^. Because the temporal specificity of the rHFOs may indicate a distinct input from the baseline HFO source, we expanded our analysis to capture changes in HFO characteristics over 10 seconds on either side of the EDs, excluding the rHFOs. We found no effect of time on power (p = 0.8300), frequency (p = 0.9178), duration (p = 0.5618), or maximum amplitude (p = 0.8219).

We next quantified differences in HFO vs rHFO characteristics within the 10 second time window. We discovered a modest but significantly higher RMS amplitude among the rHFOs vs HFOs (β = 3.1775 uV, 95% CI [1.6075, 4.7476], t(2700) = 3.9684, p<0.0001; mean 59.72 +/-13.83 vs 56.06 +/-18.58 uV, Fig. 2h). Both peak frequency (β = 20.589 Hz, 95% CI [18.29, 22.889], t(2700) = 17.556, p<0.0001; mean 162.40 +/-13.51 vs 147.37 +/-24.91 Hz, Fig. 2i) and duration (β = 11.04 ms, 95% CI [10.13, 11.96], t(2700) = 23.689; p<0.0001; mean 37.3 +/-13.1 vs 26.2 +/-6.2 ms) were greater in the rHFO group (Fig. 2j). Similarly, rHFOs exhibited a greater maximum amplitude compared to HFOs (β = 12.263 uV, 95% CI [9.1279, 15.397], t(2700) = 7.6705, p<0.0001, mean 110.25 +/-30.65 vs 96.52 +/-32.34 uV).

Interestingly, ED amplitude had a significant relationship with rHFO probability (β = −0.00113 ± 0.00014, Wald z = −8.04, p<0.001; Sup. 3). However, the odds ratio was very low (OR = 0.9989) indicating this may simply be a random effect resulting from the natural spread of the ED amplitudes (Fig.1h).

## Discussion

We discovered EDs occurring across all layers of the granular retrosplenial cortex, a region known to play a large role in memory consolidation^3,4,10,25–28^

A subset of these pathological events was associated with a unique distribution of HFOs (20-100 ms) following EDs. These rHFOs exhibited a negative relationship with time from ED onset and multiple metrics, indicating modulation by the EDs. Additionally, rHFOs were larger across all metrics than HFOs, suggesting differences in population recruitment.

Chronic implantation of electrodes has been shown to increase the potential for hippocampal spikes in the local field potential and lower the threshold for kindling in epilepsy studies^28,29^, likely via glial cell activation and neuroinflammatory processes. Neuroinflammatory markers are known to change as a result of prenatal alcohol exposure^30,31^, therefore, we suspected our TTAE group was experiencing a neuroimmune response to the electrode itself. However, upon investigation using a chronic EEG skull electrode array, we observed a similarly high number of ED-like waveforms as we observed in brain penetrating electrodes (Fig. 1b; grey triangles). Thus, while neuroinflammation may explain the small number of EDs seen in control animals (Fig. 1b-c), The TTAE group appears to be experiencing EDs via a treatment-related mechanism.

The granular retrosplenial cortex receives direct and indirect inputs from both the hippocampus and subiculum^5,32^ as well as more distant regions like the entorhinal cortex and thalamus^33^. All these regions are known to be damaged by prenatal alcohol exposure^3,34–37^. Given the EEG recording shows EDs are generalized and not focal events, it is possible polarity differences (Fig. 1f-h) may be the result of multiple circuit pathways between these interconnected regions. It is also possible rHFOs are triggered via a non-hippocampal mechanism given that only a fraction of RSC ripples are thought to be of hippocampal origin^5^. To begin untangling circuit pathways, we are currently working on building conceptual models that can be tested in silico, but more experimental work will likely be needed to further dissect where EDs originate and propagate in our TTAE mouse model.

HFOs are well established as essential to proper memory function^38–47^, therefore, we investigated how EDs may influence HFO properties in the retrosplenial cortex. The distribution of HFO timestamps revealed a large number occurring 20-100 ms after the ED onset (rHFOs; Fig. 2a-b). Interestingly, time from ED onset modulated rHFO frequency, duration, power and maximum amplitude–though this relationship did not extend to HFOs (Fig. 2c-f). EDs result from a wide variety of neurological changes, including neurodegeneration, brain injury and various forms of epilepsy. It is therefore likely the effects of EDs will differ as the specific network dysfunction differs between conditions. Gelinas^19^ et al. produced a compelling argument for the reduction in ripples following EDs in temporal lobe epilepsy to be the result of hippocampal-cortical communication being shut down via an ED-induced DOWN state in the cortex. Contrary to studies such as this, we did not observe a decrease in HFOs following EDs. However, it is plausible the rHFOs are the result of a similar pathway with the opposite effect in the retrosplenial cortex; circuit dynamics in place to generate cortical HFOs are being potentiated instead of silenced through the effects of the EDs, the product of which likely lacks the organization and thus, the information content, of non-ED-induced HFOs.

This theory is further supported by our results showing rHFOs are larger in frequency, duration, power and maximum amplitude compared to HFOs (Fig.2g-j). Studies have indicated the information content of hippocampal ripples relies on the population of cells being recruited^39,48^ and even the pattern in which they are recruited to the ripple^49–52^. We therefore hypothesize that these rHFOs are a form of pathological ripple induced by a pathological trigger (ED) and contain scrambled information due to aberrant population recruitment. We are actively investigating this relationship.

Our previous work in first and second trimester alcohol exposure revealed the pattern of maternal alcohol consumption had sex-specific effects on behavior outcomes for the offspring^53^ which supports the hypothesis that timing of exposure interacting with the phase of neurodevelopment contributes to the spectrum of outcomes seen in humans. The alcohol exposure model in the current study is performed on postnatal day 7. It is unclear if postnatal day 7 relates to a particular phase of development in the brain that lends itself to developing EDs and exposure day should be investigated in future studies.

Clinical research has shown elevated rates of abnormal EEG and epilepsy in populations with FASD, though preclinical investigation of this phenomenon is lacking. Individuals with FASD frequently cite differences in memory as having an enormous impact on their daily lives. Memory changes in epilepsy and the direct negative effect of EDs on memory are well established. We therefore present this data as a potential mechanism contributing to memory differences following TTAE through the generation of pathological rHFOs where normal HFOs should be, though future studies investigating the impact of EDs on active learning and memory consolidation are needed.

