## Supplementary figures and images for "Epileptiform Discharges Drive Unique High-frequency Oscillations Within the Retrosplenial Cortex of Mice with Third Trimester Alcohol Exposure"

### Supplementary_Figure_1

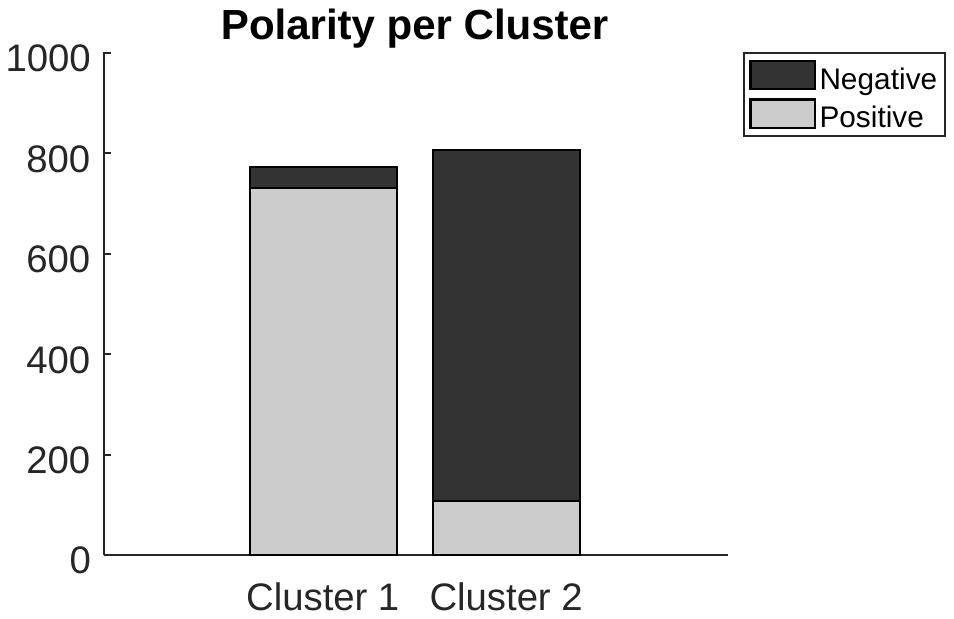

**Sup. 1; Polarity distribution between clusters**

### Supplementary_Figure_2

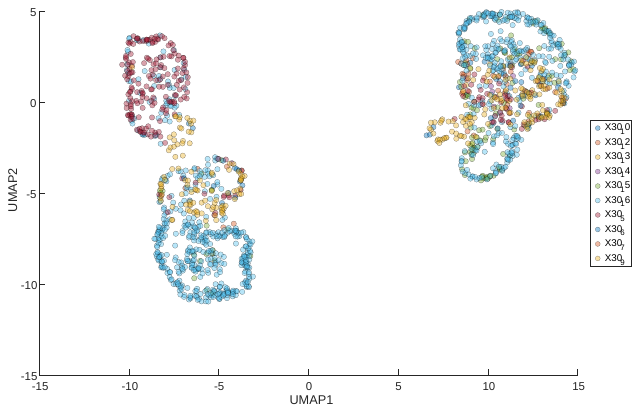

**Sup. 2; Individual animals express both cluster types**
