## Supplementary_Figure_3 for "Epileptiform Discharges Drive Unique High-frequency Oscillations Within the Retrosplenial Cortex of Mice with Third Trimester Alcohol Exposure"

**Fig. 3 ED amplitude negatively correlated with rHFO occurrence**

a) Distribution of rHFO absolute amplitude.

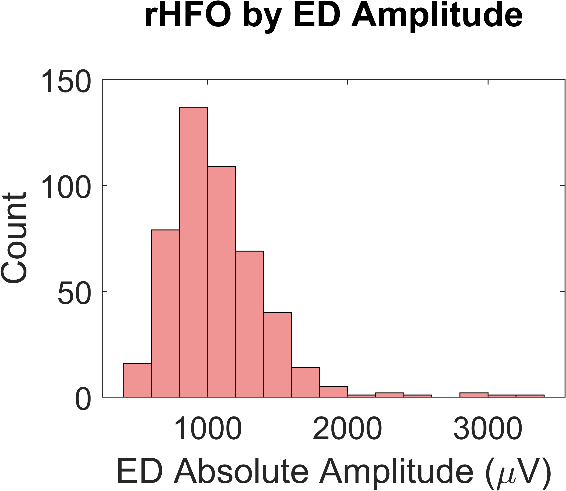

**Fig.3**

**a)**
